# Consistent thrips biological control by *Orius insidiosus* on cucumber plants despite different LED lighting regimes

**DOI:** 10.1101/2025.06.17.660193

**Authors:** Morgane L. Canovas, Adam Barrada, Paul K. Abram, Jean-François Cormier, Tigran Galstian, Martine Dorais

## Abstract

Artificial lighting, particularly light-emitting diodes (LEDs), is widely used in controlled-environment agriculture (CEA) to enhance crop productivity. Yet, light also acts as a sensory cue, directly and indirectly affecting insect pests and their natural enemies. Despite growing interest, the impacts of LEDs on multitrophic interactions remain poorly understood. This study assessed how spectral photoperiod extension modulates top-down effects (predator effects on pest densities, pest-related damage to plants, and plant defense-related gene expression) and bottom-up effects (plant morphology, light assimilation, and defense as well as growth hormone-related gene expression) in a tri-trophic system involving a plant, *Cucumis sativus* (greenhouse cucumber), a pest insect, *Frankliniella occidentalis* (western flower thrips), and a predator used for biological control, *Orius insidiosus*. Plants were exposed for two weeks to a 12-hour artificial sunlight simulating a cloudy winter day, with or without an 8-hour extension of blue, blue–red, or blue–green–red LEDs. *Orius insidiosus* reduced thrips populations by 96% ± 9%, regardless of light treatment, and decreased foliar damage due to thrips by 46% on average. Predators produced offspring under all lighting conditions, although total numbers of *O. insidiosus* were lower under blue–green–red lighting. Pest densities remained similar across all light treatments, suggesting that light supplementation did not benefit *F. occidentalis*. Blue and blue–red spectra improved plant light assimilation, and photoperiod extension generally increased dry biomass (8% ± 11%). Defense-related phytohormones were only marginally affected; surprisingly, some jasmonic acid pathway-related genes were upregulated in the presence of predators, suggesting additional plant-mediated effects. Our results underscore the potential of *O. insidiosus* to provide biological pest control services under artificial lighting regimes already used in crop production.

**Key Message:**

- Lighting in agriculture alters dynamics of pest and beneficial insects
- We asked how LED spectra influence net biological control outcomes in a tri-trophic system
- *O. insidiosus* reduced pest numbers by 96% and foliar damage by 46%
- Predators increased the expression of some jasmonic acid-related genes
- *O*. *insidiosus* showed potential for pest suppression under different lighting regimes

## Introduction

In plant–arthropod systems, light is more than a growth factor for plants —it is a sensory cue that can influence behavioral, physiological, and trophic interactions (Ben-Yakir 2020; Lazzarin et al. 2021). This is particularly relevant in controlled-environment agriculture (CEA—i.e., greenhouses and indoor farming facilities), where artificial lighting is widely used to support year-round crop production. Among available technologies, light-emitting diodes (LEDs) offer high energy efficiency and spectral flexibility, enabling precise control of plant development in CEA. LEDs are now routinely used in commercial production to enhance seedling vigor (Zhang et al. 2024), regulate ornamental flowering (Munyanont et al. 2024), and increase fruiting vegetable yields (Verma et al. 2024). While these advances offer clear agronomic benefits, they also raise ecological questions regarding the broader consequences of artificial light on other organisms inhabiting CEA systems, particularly herbivorous pests and beneficials such as introduced predators and parasitoids involved in biological control (Vänninen et al. 2010; Lazzarin et al. 2021).

Arthropod pests and beneficials rely on light cues for navigation, circadian regulation, foraging, and reproduction (Johansen et al. 2011; Shimoda and Honda 2013; Abrieux et al. 2020). Recent studies have begun to explore how specific LED spectra influence pest populations, beneficial performance and the strength of trophic interactions in horticultural contexts (Cochard et al. 2019; Park et al. 2022; Gonzalez et al. 2023; Meijer et al. 2023, 2024; Athanasiadou et al. 2024; Fraser et al. 2024; Zhu et al. 2024a; Canovas et al. 2025a, b; Savi et al. 2025). These studies, which have been conducted on a small subset of species present in CEA, suggest that shifts in light conditions could alter insect behavior, influence the dynamics of pest outbreaks and the overall effectiveness of biological control. These observed ecological responses to lighting arise through multiple mechanisms operating at different biological levels: either directly, through changes in arthropod perception and activity patterns (Johansen et al. 2011), or indirectly, *via* light-induced alterations in plant traits that affect herbivore performance and the efficacy of beneficials (Vänninen et al. 2010).

This dual pathway of lighting’s ecological effects also aligns with the trophic control framework proposed by Hunter and Price (1992), in which top-down effects refer to the regulatory influence of higher trophic levels—such as predators and parasitoids—on herbivore populations and plant damage, as well as the impact of herbivores on plants. In contrast, bottom-up effects encompass how plant quality and resource availability influence herbivores and their natural enemies. In CEA systems, artificial lighting simultaneously influences both top-down and bottom-up processes. For example, longer photoperiods under LEDs can enhance predation, parasitism, and reproduction in beneficial arthropods, thereby potentially improving pest suppression (Cochard et al. 2019; Joschinski et al. 2019; Gonzalez et al. 2023; Canovas et al. 2025b). On the other hand, LED exposure can also alter plant morphology, leaf structure, and nutritional quality, which reduce pest densities through bottom-up effects (Zhu et al. 2024a; Savi et al. 2025).

Among the plant traits influenced by light, defense-related hormonal signaling—particularly the induction of jasmonic acid (JA), salicylic acid (SA), and ethylene (ET) pathways in response to herbivore feeding—plays a central role in mediating bottom-up effects (Bürger and Chory 2019; Lazzarin et al. 2021). JA is typically induced by chewing or cell-content- feeding herbivores and orchestrates defense responses including the production of proteinase inhibitors, oxidative enzymes, and secondary metabolites. Increased JA levels have been shown to impair pests’ fitness and reduce plant damage (Abe et al. 2009; Sarde et al. 2019). SA is mainly induced by phloem-feeders or pathogens but can also contribute to herbivore-induced volatile emission and other indirect defenses, while ET modulates both JA and SA pathways (Lazzarin et al. 2021). Notably, light spectrum can significantly influence not only the production and balance of defense-related phytohormones but also plant growth-regulating hormones like auxin (AUX), which orchestrate plant development and morphology (Mirzahosseini et al. 2020; Lazzarin et al. 2021; Pierik and Ballaré 2021). Thus, plants growing under different spectra could differ in their morphology (Table 1 numb.1), their susceptibility to pests (Table 1 numb.7 & 5), and their ability to support or deter natural enemies (Table 1 numb.7, 8 & 9).

**Table 1.** Detailed description of possible top-down and bottom-up effects in the tri-trophic system in this study (*Cucumis sativus*, *Frankliniella occidentalis*, *Orius insidiosus*). Each effect (numbered 1–9) corresponds to interactions shown in the tri-trophic schematic shown in Fig. 1.

| Effect type | Description | References |
| --- | --- | --- |
| **Direct AL** | **1.** AL modulates plant energy assimilation for photosynthesis, orchestrates developmental processes such as flowering and morphology, and regulates the growth–defense trade-off. | Bantis et al. 2018<br>Pierik and Ballaré 2021<br>Lazzarin et al. 2021 |
|  | **2.** AL affects pest navigation, circadian regulation, and activity patterns, leading to changes in visual attraction, foraging behavior, population density, and survival. The visual sensitivity of *F. occidentalis* has been well characterized, and red light has notably been shown to reduce its development and feeding on plants. | Matteson et al. 1992<br>Johansen et al. 2011<br>Murata et al. 2017, 2018<br>Lopez-Reyes et al. 2022<br>Zhu et al. 2024a |
|  | **3.** AL also modulates the perception and biological rhythms of beneficials. In *Orius* spp., photoperiods of at least 10–12 hours prevent diapause, and activity, orientation, and reproduction are influenced by light spectrum and intensity. In *O. insidiosus*, spectral sensitivity varies across life stages, with photoperiod extension using blue light promoting fertility, fecundity, and adult numbers. | Henaut et al. 1999<br>Legarrea et al. 2012<br>Wang et al. 2013<br>Ogino et al. 2015, 2016, 2020<br>Labbé and McCreary 2020<br>Park et al. 2023<br>Canovas et al. 2025b |
| **Top-down** | **4.** *Orius* spp. are efficient thrips predators, often killing more prey than they consume. A 10% blue – 90% red spectrum was previously identified as optimal for predation by *O. insidiosus* on *F. occidentalis*. | Cox et al. 2006<br>Van Lenteren 2012<br>Van Lenteren et al. 2018<br>Canovas et al. 2025a |
|  | **5.** *F. occidentalis* causes plant damage through feeding and oviposition and acts as a virus vector. It significantly reduces plant photosynthesis, reproduction, and market value through fruit scarring and deformation. Cucumbers are particularly sensitive, with infestations lowering growth, photosynthetic rates, and commercial quality. | Bournier 1983<br>Hao et al. 2002 |
|  | **6.** *Orius* spp. reduce plant stress through predation, but plants may also benefit from indirect non-consumptive effects that impair thrips fitness. Notably, *O. insidiosus* maintains high performance as a predator under various AL conditions, even in complex plant environments. |  |
| **Bottom-up** | **7.** Changes in plant morphology, leaf structure, defense expression, volatile emissions, and nutrient content can influence herbivore performance and plant attractiveness. Light spectrum modulation can enhance jasmonic acid (JA)-related defenses, reducing *F. occidentalis* density and damage. JA | Abe et al. 2008, 2009<br>Escobar-Bravo et al. 2018, 2021 |
|  | induction by thrips feeding has been linked to lower oviposition and larval development. Optimized light intensity can also increase thrips resistance through greater trichome density and allelochemical production. |  |
|  | **8.** AL indirectly affects predator performance by modifying prey availability and quality. Changes in thrips density, size, or nutritional value under different light conditions could influence *O. insidiosus* feeding behavior and reproductive success. | Vänninen et al. 2010<br>Johansen et al. 2011 |
|  | **9.** Structural changes in plants induced by AL—such as increased trichome density—can influence predator mobility, access to prey, and overall efficiency. Additionally, light-driven shifts in the emission of volatile organic compounds (VOCs), like terpenes, can modulate predator behavior by either attracting or repelling beneficial arthropods. | Vänninen et al. 2010<br>Lazzarin et al. 2021 |

While several studies have independently addressed top-down or bottom-up responses to LED lighting on single trophic levels (Vänninen et al. 2010; Lazzarin et al. 2021), the interacting effects of these pathways within whole plant–pest–beneficial tri-trophic systems remain largely unexplored. Current evidence comes from only a few model systems, but does suggest that LED lighting can lead to similar or improved pest management and, in some cases, enhanced plant agronomic traits and defense responses (Meijer et al. 2023, 2024; Fraser et al. 2024; Zhu et al. 2024a; Savi et al. 2025). More studies are needed to determine how artificial lighting-induced changes propagate across trophic levels and ultimately influence the net amount of plant protection provided by biological control and plant resistance.

In this study, we examined the impact of spectral photoperiod extension on ecological dynamics within a tri-trophic model system comprising cucumber (*Cucumis sativus*), a vegetable crop widely grown under artificial lighting (Dyśko and Kaniszewski 2021), *Frankliniella occidentalis*, a major pest (Reitz et al. 2020), and *Orius insidiosus*, a generalist predatory bug commonly introduced in CEA (Van Lenteren et al. 2018). Selected for its agronomic relevance and sensitivity to different lighting regimes (Hao et al. 2002; Garcia and Lopez 2020; Lopez-Reyes et al. 2022; Canovas et al. 2025b, a), this tri-trophic system allowed us to investigate some of the top-down effects on thrips control and plant damage through predator and pest responses, as well as a subset of bottom-up effects on pests and predators *via* light-induced changes in plant morphology and hormone signaling (Fig.1).

**Fig. 1.**
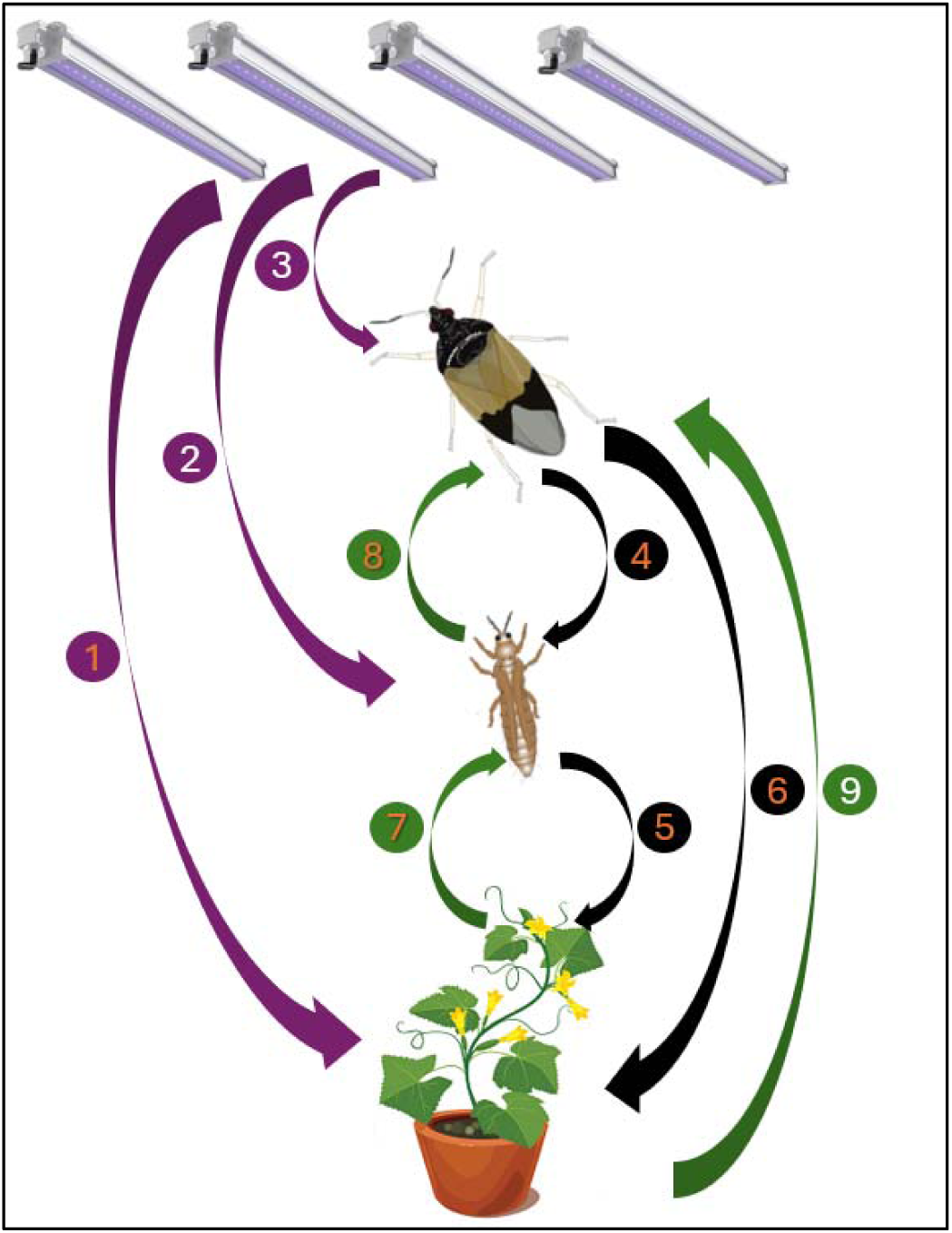
Schematic representation of tri-trophic interactions in the model plant – pest – predator system (*Cucumis sativus* – *Frankliniella occidentalis* – *Orius insidiosus*), including the influence of artificial lighting (AL). Purple, black, and green arrows represent direct AL effects, top-down, and bottom-up interactions, respectively. Interactions investigated in the present study are numbered in orange. See Table 1 for a detailed description of interactions numbered 1–9, along with the corresponding references. Direct effects of AL include its influence on plant photosynthesis, growth, and defense (1), on pest behavior, abundance, and survival (2), and on predator behavior and population dynamics (3). Top- down interactions include the ability of *Orius* spp. to effectively control thrips populations, with certain light spectra enhancing this predatory activity (4), the detrimental effects of *F. occidentalis* on plant growth and fruit quality (5), and the mitigation of plant stress under AL through predator presence (6). Bottom-up effects reflect the role of lighting in altering plant morphology and defense traits (7), its indirect effects on prey quality and quantity *via* plant mediation (8), and the modulation of predator behavior through plant traits influenced by AL (9)

Using a programmable horticultural lighting system, we exposed plants, pests, and predators to a 12-hour baseline solar light regime simulating a cloudy winter day, with or without an 8- hour photoperiod extension. The extension used one of three spectral treatments reported to support *O. insidiosus* predation on thrips (Canovas et al., 2025a) and developmental performance (Canovas et al., 2025b): blue (B), blue–red (BR), or blue–green–red (BGR).

After two weeks, we monitored predator and pest populations, visible damage, light assimilation, plant morphology and growth- and defense-related foliar phytohormones expression.

In terms of top-down effects, based on previous observations of *O. insidiosus*, we expected that blue light supplementation would increase predator fecundity (Labbé and McCreary 2020; Canovas et al. 2025b), while BR light would enhance predation behavior (Canovas et al. 2024), both contributing to stronger pest suppression and reducing plant damage.

Additionally, we predicted that BR lighting may independently suppress *F. occidentalis* development and feeding activity (Murata et al. 2017, 2018; Zhu et al. 2024a). Regarding bottom-up effects, we predicted that B and BR spectra—aligned with chlorophyll absorption peaks—would enhance light assimilation and plant biomass (Bantis et al. 2018), potentially favoring higher pest populations in the absence of natural enemies. Thrips feeding in predator-free conditions was anticipated to trigger defense-related phytohormones, especially JA (Abe et al. 2008), potentially reducing pest oviposition and larval development (Abe et al. 2009). In contrast, predator presence might mitigate stress-related hormonal responses regardless of lighting. JA signaling was expected to be most strongly induced by pest feeding under BGR light, while BR light was predicted to increase SA and ET-related gene expression (Mirzahosseini et al. 2020). Finally, BR was assumed to induce the expression of AUX-related genes(Liu et al. 2011; Guo et al. 2016). By integrating plant physiology, pest performance, and predator responses, we aimed to inform lighting strategies that enhance both biological control and crop resilience in CEA systems.

## Material and methods

### Lighting device and treatments

To assess the top-down and bottom-up effects of both photoperiod extension and light spectrum on our tri-trophic system, we employed a laboratory setup similar to Canovas et al. (2025a, b). The experiment was conducted in laboratory, in four opaque growth tents (©2022 VIVOSUN, Philadelphia, Ontario; dimensions: 121.9 cm × 121.9 cm × 182.9 cm), each equipped with dynamic spectrum horticultural LED systems (©2022 SOLLUM Technologies, Montreal, Quebec; model SF05-A). These LEDs allowed for real-time adjustments of both spectral composition and light intensity through the SUN as a Service® platform (SUNaaS). Environmental conditions were controlled throughout the experiment, maintaining a temperature of 21 ± 2°C and relative humidity of 60 ± 2%, regulated using a combination of climate control devices (©2022 Inkbird, Shenzhen, China; models ITC-308 and IHC-200; ©2022 VIVOSUN, Philadelphia, Ontario, CA; model 43237-2; ©2022 ALACRIS, Ottawa, Ontario, CA; 4L humidifier).

Each tent was assigned to one of the tested light treatments. Based on prior findings (Canovas et al. 2025a, b), we selected three light spectra commonly used in horticulture and known to support *O. insidiosus* development and predation on thrips: 100% Blue (B), 25% Blue – 25% Green – 50% Red (BR), and 10% Blue – 90% Red (BGR). These spectral treatments were applied as supplemental lighting for 8 hours after a simulated 12-hour of low-light natural sunlight photoperiod, mimicking winter greenhouse conditions typical of northern latitudes (Fig. 2). The "natural sunlight" 12-hour light cycle was designed based on average spectrometric data for the Quebec City region provided by the World Meteorological Organization (WMO) and integrated *via* the SUNaaS platform, including a sunrise and sunset phase (Honnorine Lefevre, agr., personal communication; SOLLUM Technologies). A control treatment (CTRL), without photoperiod extension, was included for comparison. Light intensity during the "natural sunlight" phase was standardized at 240 µmol/m²/s, with approximately 30 µmol/m²/s during the extension phase. This range was selected based on previous observations indicating no significant changes in *O. insidiosus* predation behavior within these light intensity parameters (Canovas et al., 2025a). Therefore, the daily light integral (DLI) was 1123 mol/m²/day, except for the control treatment (10,37 mol/m²/day).

**Fig. 2.**
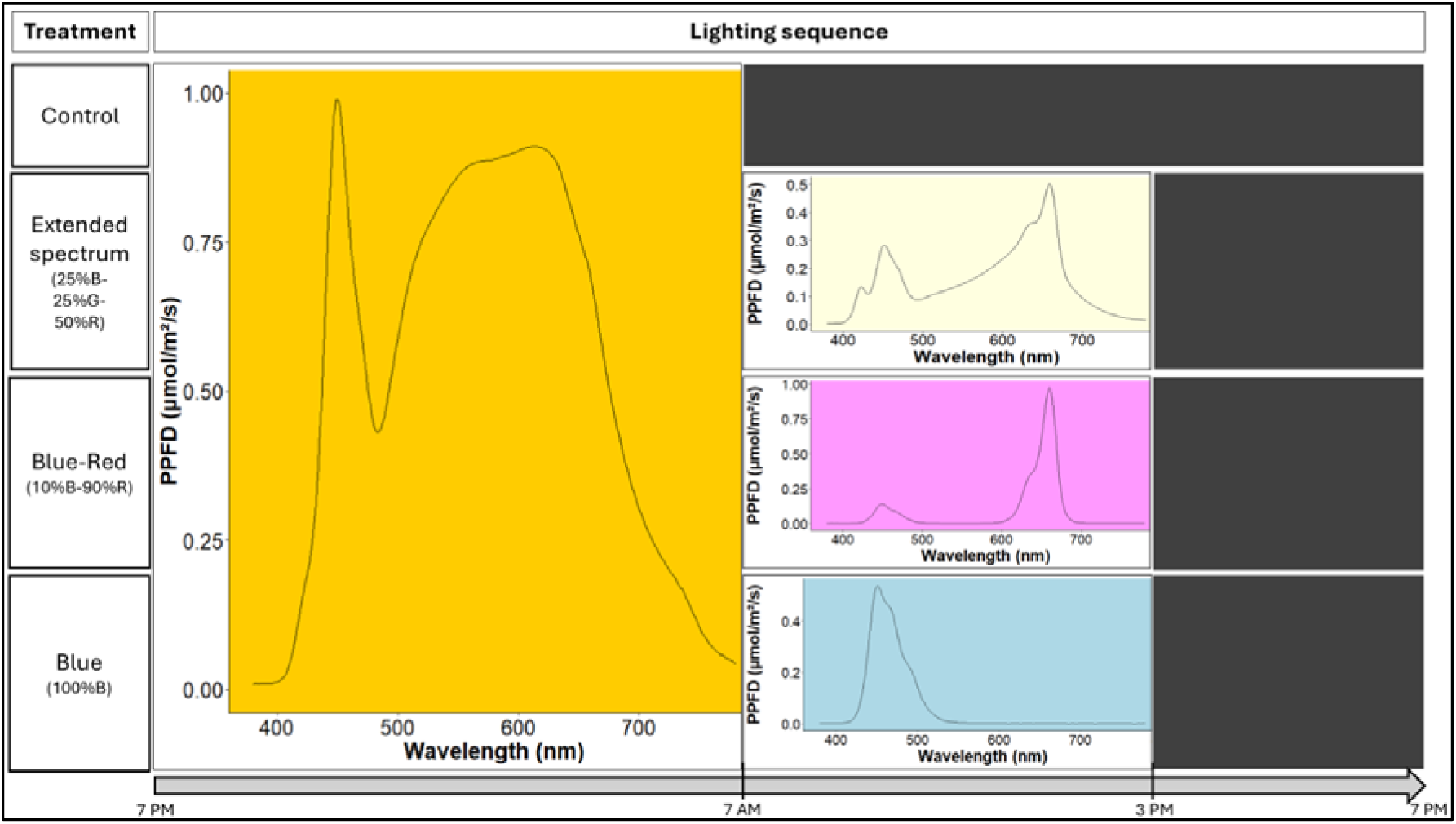
Lighting sequences tested in growth tents during the cage experiment. The 24-hour cycle was structured in three phases: an initial exposure to a simulated low intensity solar spectrum (shown in orange), followed by a photoperiod extension using one of the three tested light spectra (represented in beige, violet, or blue, depending on the treatment; absent in the control), and concluding with a dark phase (shown in black)

To distinguish top-down from bottom-up effects induced by light, during the experiment three insect treatments were nested within each artificial lighting sequence tested: 1) cucumber plant only, 2) cucumber plant with pest thrips, and 3) cucumber plant with pest thrips and predatory bugs. The experiment followed a full factorial design, testing four light treatments × three insect treatments, with three biological replicates per combination, repeated across four independent temporal blocks. This resulted in 12 independent plant observations per combination, totaling 36 observations per light treatment and 48 observations per insect treatment.

### Plant cultivation and insect rearing

*Cucumis sativus* (Marketmore70 variety, organic seeds; ©NORSECO, Laval, QC, Canada) were sown in standard 5 ½- inch pots filled with a commercial growing mix (PRO-MIX BX; PremierTech Biotechnologies, Rivière-du-Loup, QC, Canada). Seedlings were immediately exposed to one of the tested light treatments, with nine plants per treatment. Progressive weekly fertilization with a 20-20-20 (N-P-K) fertilizer solution (Plant-Prod 20-20-20 Classic; PlantProducts, Leamington, ON, Canada) began at true-leaf emergence at 100ppm (in week 2) and gradually increased to 200ppm by weeks 3-6.

*Frankliniella occidentalis* were sourced from a permanent colony maintained at Laval University in a separate growth chamber from the light trials (CONVIRON, Winnipeg, MB, Canada; 25 ± 2°C, 65 ± 2% RH, photoperiod 16:8 h light:dark). New individuals were regularly introduced from university greenhouse complexes or commercial garden centers, with species identity previously verified under a stereomicroscope (OLYMPUS SZ61, Quebec, QC, Canada) (Mound and Kibby 1998). Rearing was conducted in transparent 1 L Mason jars with Nitex (160 µm nylon mesh) lids. Each jar contained ∼2 cm of vermiculite and six commercial green bean pods, serving as both food and oviposition substrates. Pods were disinfected for 10 minutes in a 5% bleach solution (LAVO PRO 6) before introduction, following Labbé et al. (2018). New pods were added weekly. To mimic thrips infestation in CEA context, six out of nine cucumber plants per light treatment were individually placed in thrips-proof cages (BugDorm, Taichung, Taiwan; model 4E2260) and inoculated with fifteen female thrips of similar size and color, allowing for both population establishment and acclimation to the tested light sequence for 10 days prior to the experiment (Fig. 3). By day four, a new generation of thrips is expected to hatch and begin feeding, contributing to damage (Steenbergen et al. 2020).

**Fig. 3.**
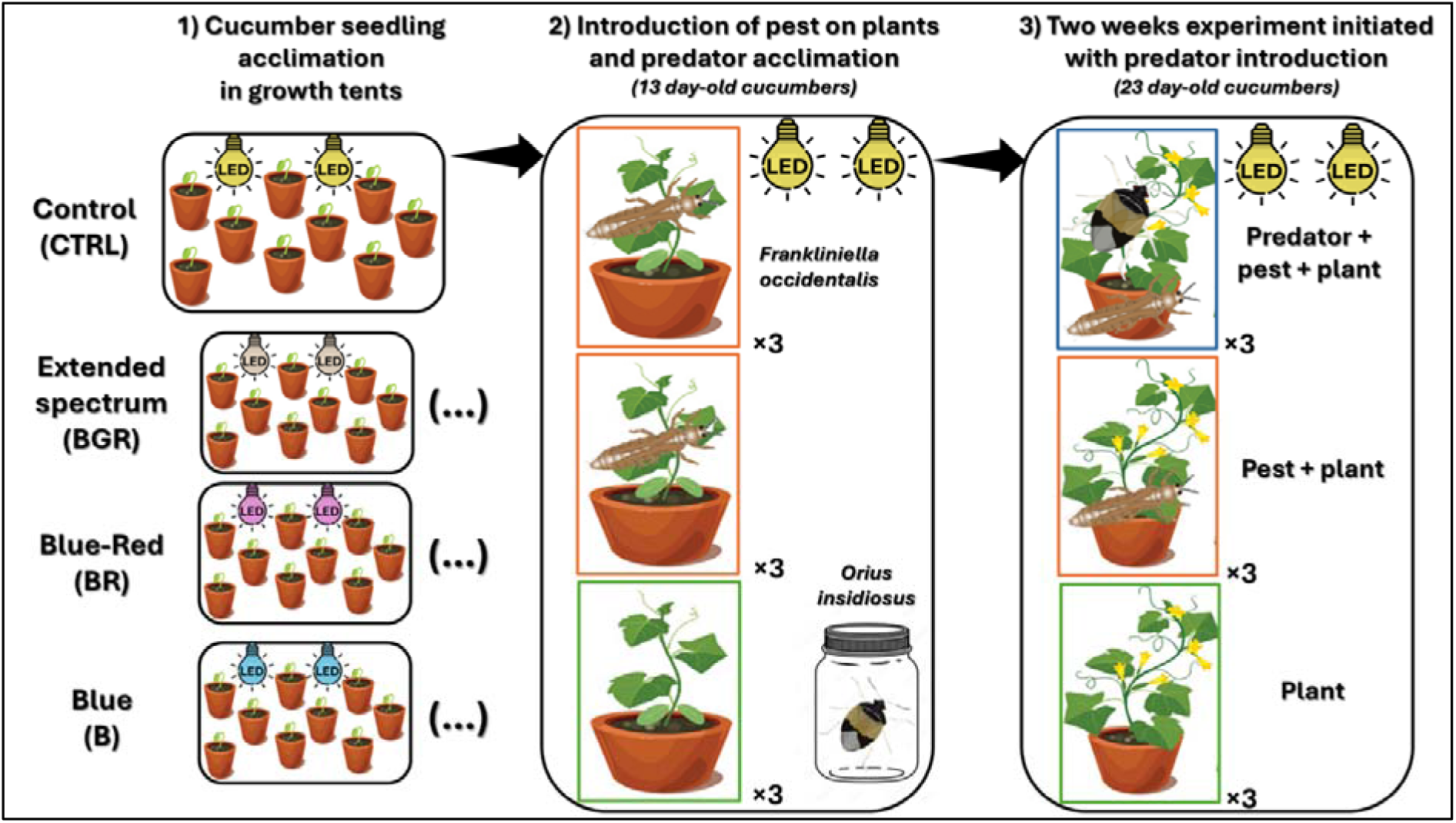
Simplified protocol for one temporal block of the cage experiment involving cucumber plants (*Cucumis sativus*), pest thrips (*Frankliniella occidentalis*) and nymph and adult female predatory bugs (*Orius insidiosus*)

*Orius insidiosus* adults (2–3 days old) were obtained from ©ANATIS Bioprotection (Saint-Jacques-le-Mineur, QC, Canada) for periodic rearing during the trials. Upon receipt, 500 individuals were housed in 1 L Mason jars with Nitex lids. Jars contained a mixture of buckwheat husks and vermiculite (∼3 – 4 cm) along with absorbent paper to create structural complexity, reducing cannibalism and maintaining humidity (Schmidt et al. 1995). Six disinfected green bean pods (as described for *F. occidentalis*) served as water and oviposition substrates. As a food source, six 2 cm diameter stickers coated with *Ephestia kuehniella* eggs (©ANATIS Bioprotection) were provided. Bean pods and egg sources were replaced every three days, following Labbé et al. (2018). To mimic predator introduction by growers in CEA context, a Mason jar was placed in each light treatment tent upon predators’ receipt for a 10- day acclimation period prior to the experiment (Fig. 3).

### Experimental protocol

We aimed to assess the effects of both photoperiod extension and light spectrum on our tri- trophic system through top-down and bottom-up processes, using 23-day-old cucumber plants along with pest and predatory insects previously acclimated to the tested light sequences (see above). On the day of the beginning of the experiment, four adult females and four 2^nd^ instar nymphs *O. insidiosus* were introduced into three of the six cages previously inoculated with thrips (Doğramaci et al. 2011; Labbé et al. 2018), while the remaining three cages served as predator-free controls (Fig. 3). The predators, aged approximately 14–17 days for the adults and 2-3 days for the nymphs (with a lifespan of ∼30 days; Schmidt et al. 1995), were individually collected and transferred into 1.5 ml sterile tubes for sex and stage confirmation under a stereomicroscope based on morphological criteria (Isenhour and Yeargan 1981). To support predator establishment and enhance longevity, *Ephestia kuehniella* (Zeller) eggs were added just before predator introduction, following Labbé et al. (2018). This practice has been shown not to reduce the control efficiency of *O. insidiosus* against *F. occidentalis* (Labbé et al., 2018). Additionally, in commercial greenhouse settings, *O. insidiosus* is often supplied mixed with a dry substrate (e.g., perlite or buckwheat husks) and *E. kuehniella* eggs, which are directly applied to crop foliage during predator introduction. The addition of eggs as a supplemental food source for predators was repeated weekly after predator introduction, coinciding with cucumber plant fertilization.

### Data collection

#### Pest and Predator Insect Densities

Fifteen days after predator introduction, each plant inoculated with insects (Plant + Pests and Plant + Pests + Predator treatments) was sampled using an insect vacuum. All life stages of prey and predators were collected separately, stored in labeled tubes, and frozen at −20°C for subsequent counting under a stereomicroscope. No insects were detected in the plant-only treatment.

#### Plant Morphological Traits

Following insect sampling, the following morphological measurements were recorded: plant height, width (maximum distance between two opposite leaf tips), diameter and number of leaves. Fresh shoot biomass was assessed after performing the measurements described in *Plant Light Assimilation* section. Dry biomass was determined after drying plant samples in at 60 °C for 5 days in an oven.

#### Plant Light Assimilation

After morphological measurements, plants were returned to their respective light treatments (B, BR, BGR spectra, or the "sunlight" spectrum for CTRL). Following at least 30 minutes of light exposure, chlorophyll content was measured using a SPAD 502 PLUS meter (Spectrum Technologies Inc., Aurora, IL, USA) on the adaxial surface of the fifth fully expanded leaf, counted from the top of the plant. The physiological health of the photosynthetic apparatus was then assessed using a Handy PEA+ fluorimeter (Hansatech Instruments, King’s Lynn, Norfolk, UK) after 20 minutes of dark adaptation, based on two parameters: the Fv/Fm ratio, reflecting the maximum quantum yield of photosystem II (an indicator of photoinhibition or photosynthetic stress; then referred to as quantum yield or QY), and the Performance Index (PI), which integrates multiple photosynthetic steps to provide a sensitive measure of overall photosynthetic capacity. For all three indicators, a mean value was calculated from three consecutive measurements per plant to account for potential instrumental error.

#### Silvering Damage Assessment

Thrips feeding on leaves causes characteristic silvering marks, with both adults and larvae contributing to the damage (Bournier 1983). Given that feeding activity predominantly occurs on the abaxial leaf surface (Visschers et al. 2018), the abaxial side of each cucumber leaf was scanned at a resolution of 5100 × 7020 pixels, and the total area of thrips-induced silvering damage on each plant was quantified using ImageJ (version 1.53t, Fiji distribution, Schindelin et al. 2012), considering lesions of 0.01 cm² and larger.

#### Plant defense and growth gene expression

After scanning for pest damage, the fourth and fifth leaves (from the bottom) of each cucumber plant were collected and flash-frozen in liquid nitrogen. Total RNA was extracted from ∼250 mg of tissue using the EZ-10 Spin Column Plant RNA Miniprep Kit (Bio Basic, Markham, Canada) and subsequently treated with RQ1 DNase (Promega, Madison, USA). RNA concentration and purity were verified spectrophotometrically, and cDNA synthesis was performed with 14–500 ng of RNA, depending on extraction yield, using random primers and oligo(dT) with the PrimeScript RT Reagent Kit (Takara Bio, Kusatsu, Japan). The samples with insufficient yield were excluded from the analysis. Quantitative PCR (qPCR) reactions (35 µL) containing SYBR™ Green I (1:20,000 final dilution; Thermo Fisher Scientific Inc., Waltham, USA), 1.75 units of Taq polymerase (NEB, Pickering, Canada), 0.2 mM dNTPs, and 0.2 µM primers (Supplementary Material 1) were run on a CFX Opus 96 Real-Time PCR System (Bio-Rad, Hercules, USA). The qPCR program included an initial denaturation at 95L°C for 30 s, followed by 45 cycles of 95L°C for 10 s, 60L°C for 30 s and 68L°C for 30 s, ending with a melting curve analysis to confirm product specificity. Cq values were adjusted using primer efficiencies determined with PCR Miner (Zhao and Fernald 2005).

ACT, UBQ, CACS, CYP and EF1 were used as reference genes. Based on previous studies in *Cucumis sativus*, primers targeting key genes involved in major plant growth and defense pathways against cell content-feeding herbivores were selected (Supplementary Material 1). Relative gene expression was calculated using the 2^−ΔΔCt method, with the CTRL × plant only treatment combination used as the reference for fold change expression. Z-scores were calculated for each combination (lighting × insect treatment), and extreme outlier observations (Z-scores > 3) were excluded to minimize the impact of experimental noise. The final number of observations after excluding samples with insufficient yield ranged from 4 to 11 per lighting × insect treatment combination.

The entire experiment was repeated four times using independent batches of predators to account for potential variability in predator vigor across shipments of *O. insidiosus* from the commercial biological control supplier. Insect–plant combinations were randomized within each light treatment, and light treatments were reassigned to different growth tents at the start of each temporal blocks to minimize positional and environmental bias.

### Statistical analysis

All analyses were conducted using R software (version 4.4.1; R Core Team, 2024), with the statistical significance threshold set at p = 0.05. Generalized linear mixed models (GLMMs) were fitted using the `glmmTMB` function from the `glmmTMB` package (Brooks et al., 2025), and model selection among candidate error distributions was performed using Akaike’s Information Criterion (AIC), computed with the base `AIC` function. Each model contained insect treatment, lighting treatment, and their interaction as fixed factors. Temporal blocks, each using distinct insect batches, were modeled as random effects to account for variability across runs, with biological replicates nested within them.

For insect-related parameters, we specified: a generalized Poisson error distribution (log link) for total predator counts; a negative binomial type 2 distribution (log link) for total thrips counts; and a gamma distribution (log link) for silvering damage area. Fresh and dry biomass, as well as leaf number, were analyzed with a Tweedie distribution (log link). Plant height, width, and diameter were modeled using a Gaussian distribution (log link). For light assimilation parameters, SPAD and PI indices were analyzed using a Tweedie distribution (log link), and QY (Fv/Fm ratio) was analyzed using a zero-inflated beta distribution (logit link). The expression of growth- and defense-related genes was analyzed using a gamma distribution (log link). Pairwise comparisons among levels of significant factors were conducted using estimated marginal means (EMMs) with Tukey adjustments for multiple comparisons (Graves et al., 2024; Lenth, 2025). p-values were derived from asymptotic Wald tests, assuming infinite denominator degrees of freedom, except for models using a Gaussian distribution. Residual diagnostics were performed using the `DHARMa` package (Hartig, 2024), with simulated residuals visually inspected to assess deviations from model assumptions, including residual uniformity, overdispersion, and zero inflation. Conditional pseudo-R² values were calculated as the squared correlation between the observed response values and the fitted values from the model including random effects. Graphical representations, obtained with package `ggplot2`, illustrate data adjusted from the full model, based on estimated marginal means (Lenth, 2025). Analyses were validated by professional statistician Gaétan Daigle, MSc (Statistical Consulting Service, Laval University).

## Results

### Insect Densities and Silvering Damage

Predator abundance was influenced by light spectrum (Table 2), with *O. insidiosus* adults and nymphs being 39% less abundant under BGR lighting compared to the CTRL condition (Fig. 4A). In contrast, pest (*F. occidentalis*) abundance was not affected by lighting treatment but was strongly reduced by *O. insidiosus*, showing an average 96% ± 9% decline in the presence of predators (Fig. 4B). Similarly, leaf damage area decreased by 46% when predators were present (Fig. 4C). A marginally significant interaction between light spectrum and insect treatment was detected for leaf damage area (p = 0.069). Under BR lighting, predators significantly reduced silvering damage, with affected area decreasing from 1.62 mm² ± 0.7 in pest-only treatments to 0.38 mm² ± 0.16 in predator treatments (Supplementary Material 2).

**Fig. 4.**
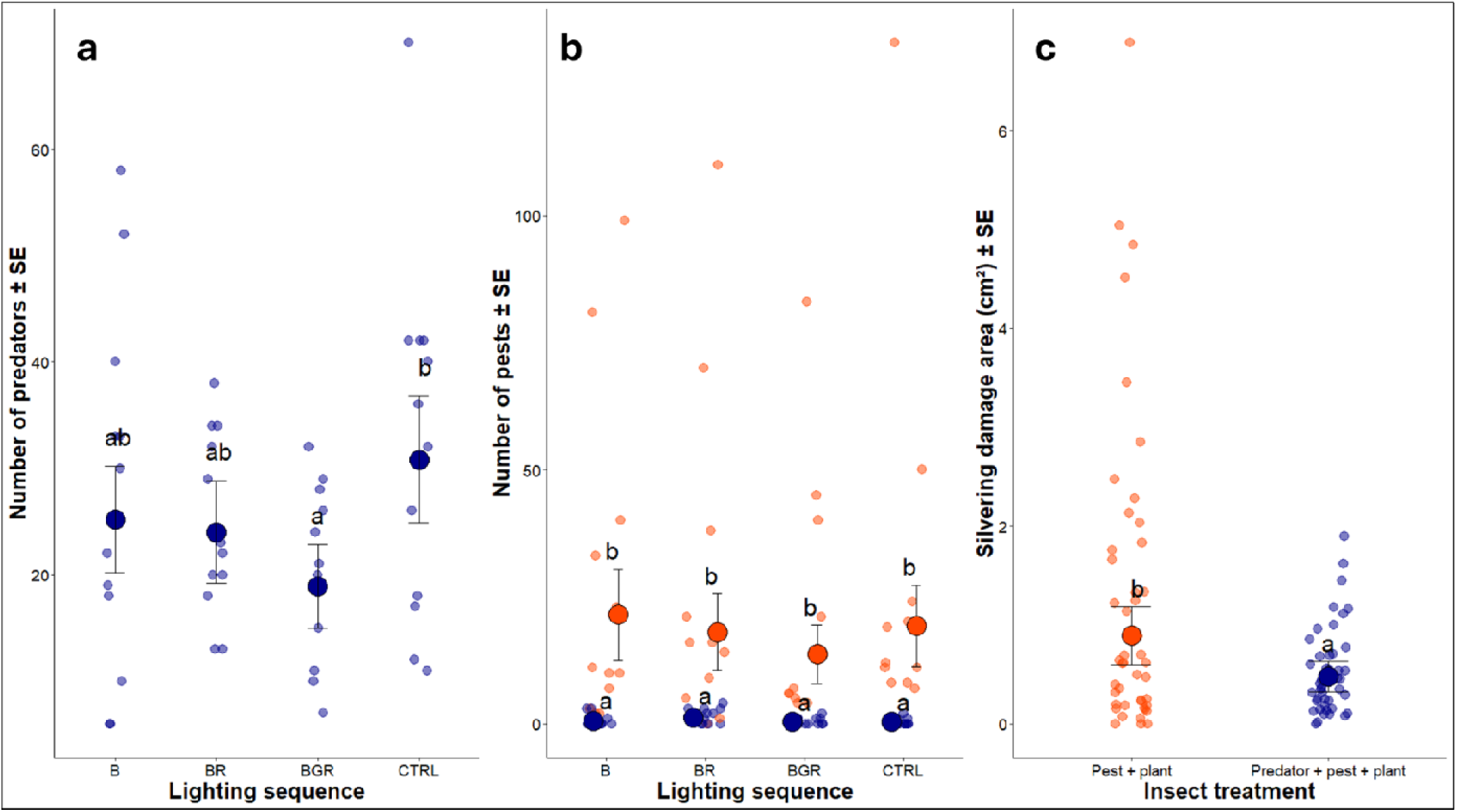
Insect-related parameters measured two weeks after exposing cucumber plants to different lighting spectra either in the absence of insects, with pests (*F. occidentalis*), or with pests and predators (*O. insidiosus*): a) Number of predators (± SE) according to lighting sequence; b) Number of pest thrips (± SE) according to the interaction between lighting sequence and insect treatment; c) Silvering damage area caused by thrips feeding (± SE) according to insect treatment. Pale points show raw data; darker points indicate model-predicted means. In panel C, the lighting sequence is not shown as it was not statistically significant (p-value = 0.368)

**Table 2.** Statistical comparison of insect-related parameters of insect combinations exposed to different lighting sequences.

| Parameter | Factor | F-value<sub>df</sub> | P-value | Conditional pseudo-R<sup>2</sup> |
| --- | --- | --- | --- | --- |
| Number of predators | spectrum | $F_{(3, \infty)} = 3.447$ | 0.016 | 0.53 |
| Number of pests | spectrum | $F_{(3, \infty)} = 1.473$ | 0.220 | 0.54 |
| | insect | $F_{(1, \infty)} = 162.514$ | <0.0001 | |
| | spectrum × insect | $F_{(3, \infty)} = 1.193$ | 0.311 | |
| Silvering damage area | spectrum | $F_{(3, \infty)} = 1.053$ | 0.368 | 0.44 |
| | insect | $F_{(1, \infty)} = 6.092$ | 0.014 | |
| | spectrum × insect | $F_{(3, \infty)} = 2.363$ | 0.069 | |

### Plant Morphological Traits

Cucumber plant biomass was affected by both insect treatment and light spectrum, but with distinct patterns (Table 3). Fresh biomass was reduced by 17.6% in pest-only treatments compared to plant-only controls, while this reduction was smaller (9.5%) when predators were also present, indicating that predator presence mitigated pest-induced loss of fresh plant biomass (Fig. 5A). Additionally, plant fresh biomass was 12% higher under B light compared to BR light (Fig. 5B). Dry plant biomass, however, was influenced solely by light treatments, with photoperiod extensions generally yielding higher dry mass (8% ± 11%); plants under control lighting were 12.9% lighter than those subjected to photoperiod extension with BGR light (Fig. 5C).

**Fig. 5.**
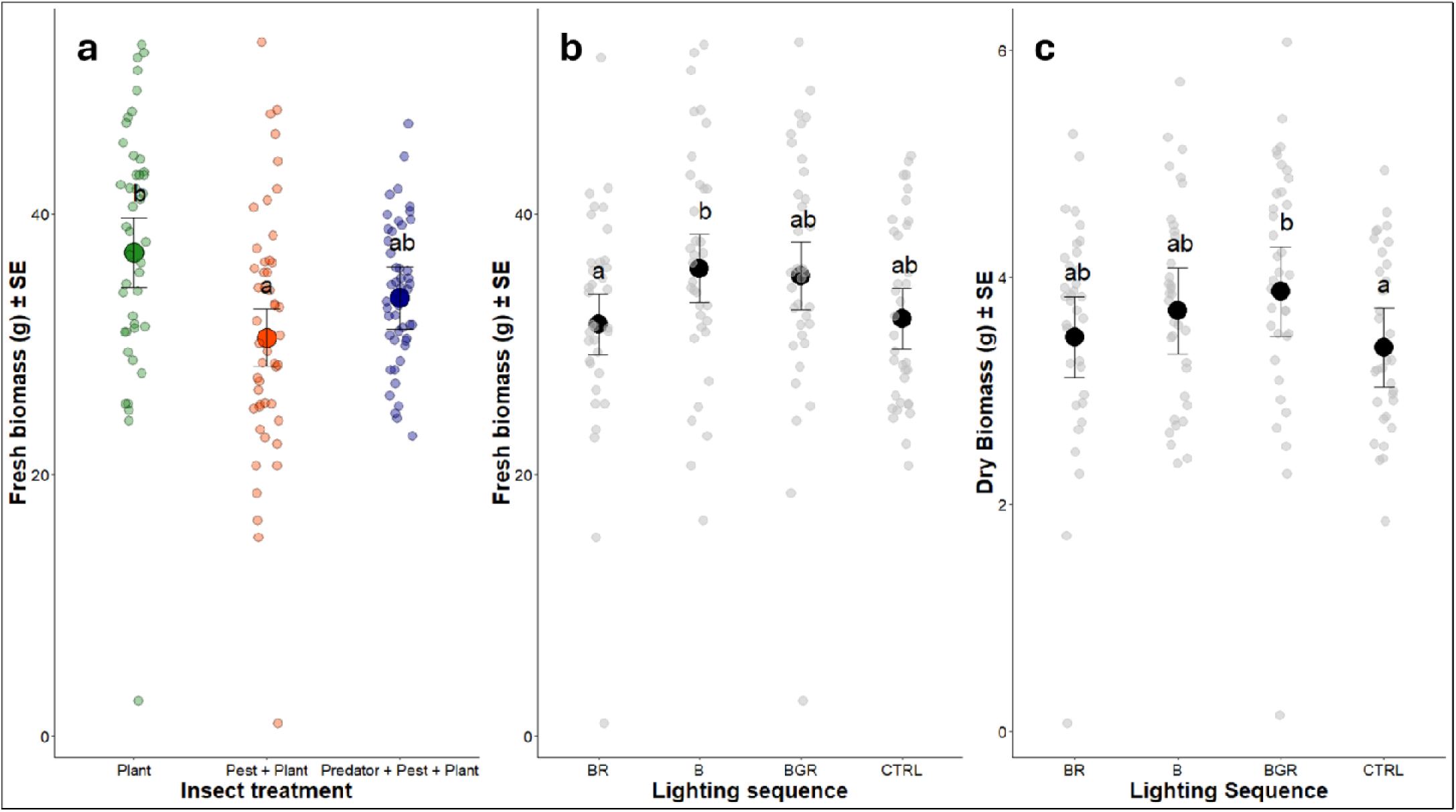
Biomass-related parameters measured two weeks after exposing cucumber plants to different lighting spectra either in the absence of insects, with pests (*F. occidentalis*), or with pests and predators (*O. insidiosus*): a) Fresh biomass (± SE) according to insect treatment; b) Fresh biomass (± SE) according to lighting sequence; c) Dry biomass (± SE) according to lighting sequence. Pale points show raw data; darker points indicate model-predicted means

**Table 3.** Statistical comparison of plant morphology- and light assimilation-related parameters of plant-insect combinations exposed to different lighting sequences.

| Parameter | Factor | F-value<sub>df</sub> | P-value | Conditional pseudo-R<sup>2</sup> |
| --- | --- | --- | --- | --- |
| **Fresh biomass** |  |  |  | 0.49 |
| | spectrum | $F_{(3, \infty)} = 3.520$ | 0.014 | |
| | insect | $F_{(2, \infty)} = 10.000$ | <0.0001 | |
| | spectrum × insect | $F_{(6, \infty)} = 1.554$ | 0.156 | |
| **Dry biomass** |  |  |  | 0.27 |
| | spectrum | $F_{(3, \infty)} = 3.592$ | 0.013 | |
| | insect | $F_{(2, \infty)} = 0.565$ | 0.569 | |
| | spectrum × insect | $F_{(6, \infty)} = 1.389$ | 0.215 | |
| **SPAD** |  |  |  | 0.44 |
| | spectrum | $F_{(3, \infty)} = 6.763$ | 0.0001 | |
| | insect | $F_{(2, \infty)} = 1.438$ | 0.238 | |
| | spectrum × insect | $F_{(6, \infty)} = 1.343$ | 0.234 | |
| **QY** |  |  |  | 0.40 |
| | spectrum | $F_{(3, \infty)} = 6.700$ | 0.0002 | |
| | insect | $F_{(2, \infty)} = 1.830$ | 0.160 | |
| | spectrum × insect | $F_{(6, \infty)} = 1.180$ | 0.314 | |
| **PI** |  |  |  | 0.34 |
| | spectrum | $F_{(3, \infty)} = 4.498$ | 0.004 | |
| | insect | $F_{(2, \infty)} = 1.686$ | 0.185 | |
| | spectrum × insect | $F_{(6, \infty)} = 1.132$ | 0.340 | |
| **Plant height** |  |  |  | 0.67 |
| | spectrum | $F_{(3, 126)} = 12.115$ | <0.0001 | |
| | insect | $F_{(2, 126)} = 38.827$ | <0.0001 | |
| | spectrum × insect | $F_{(6, 126)} = 1.291$ | 0.266 | |
| **Plant width** |  |  |  | 0.31 |
| | spectrum | $F_{(3, 125)} = 10.004$ | <0.0001 | |
| | insect | $F_{(2, 125)} = 3.642$ | 0.029 | |
| | spectrum × insect | $F_{(6, 125)} = 1.738$ | 0.118 | |
| **Plant diameter** |  |  |  | 0.63 |
| | spectrum | $F_{(3, 125)} = 2.302$ | 0.080 | |
| | insect | $F_{(2, 125)} = 1.537$ | 0.219 | |
| | spectrum × insect | $F_{(6, 125)} = 1.053$ | 0.395 | |
| **Number of leaves** |  |  |  | 0.52 |
| | spectrum | $F_{(3, \infty)} = 9.648$ | 0.022 | |
| | insect | $F_{(2, \infty)} = 2.344$ | 0.310 | |
| | spectrum × insect | $F_{(6, \infty)} = 0.276$ | 0.949 | |

### Plant Light Assimilation

The cucumber plants’ light assimilation indicators were strongly shaped by light spectra (Table 3), with B light consistently producing high values across SPAD, quantum yield (QY), and performance index (PI) measurements (Fig. 6). SPAD was 7.4% higher under BR compared to BGR lighting, QY was 1.4% higher under B compared to CTRL, and PI was 18.4% higher under BR compared to CTRL. Additional morphological traits varied with lighting treatment, while plant width and height were also influenced by insect treatment; however, no further effect of the latter was detected on overall plant diameter (Table 2; see Supplementary Material 3 for illustration).

**Fig. 6.**
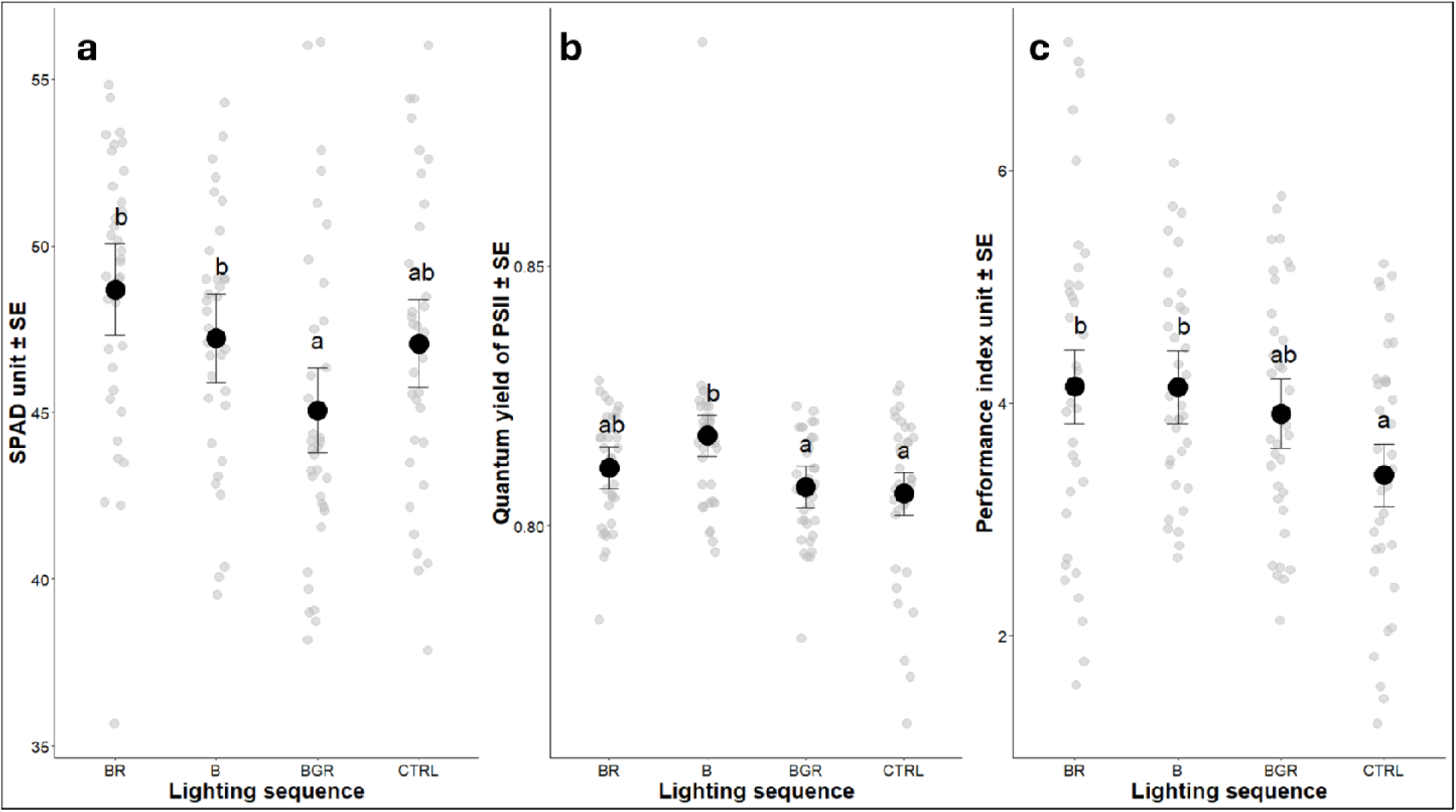
Light assimilation-related parameters measured two weeks after exposing cucumber plants to different lighting spectra either in the absence of insects, with pests (*F. occidentalis*), or with pests and predators (*O. insidiosus*) according to lighting sequence: a) SPAD unit (± SE); b) Quantum yield of PSII (± SE); c) Performance index unit (± SE). Grey points show raw data; darker points indicate model-predicted means

### Plant defense and growth gene expression

Of the 17 tested cucumber plant genes, 6 responded to the experimental treatments (Table 4). Among jasmonic acid (JA)-associated genes, *LOX1* showed a significant interaction between lighting treatment and insect treatment, with expression 2.5 times higher in the pest-only condition compared to predator-present under BR light (Fig. 7A). *MYC2* and *WRKY20* were influenced by insect treatment, showing 3- and 2-fold higher expression, respectively, in the presence of predators compared to the no-insect control and the pest-only treatment (Fig. 7B- C). *MYC2* was also influenced by lighting treatment, with 1.7-fold higher expression under BR compared to BGR light. For auxin (AUX)-associated genes, *SAUR61* was induced by insect treatment, showing a 3.5-fold increase in expression in the presence of predators compared to both the no-insect control and the pest-only treatment (Fig. 8A). A significant spectrum × insect interaction was detected for *IAA26*: under BR light, expression levels were 5 times higher in pest-only plants than in plant-only controls (Fig. 8B). Lastly, the salicylic acid (SA)-associated gene *NPR1* was independently affected by both light spectrum and insect treatment: NPR1 transcript levels wer twice as high higher in plant-only compared to pest-only plants, and 1.5-fold higher under B light compared to CTRL (Fig. 9). While these effects were statistically significant, the conditional pseudo-R² values ranged from 0.13–0.29, indicating that the models explained only a small to modest portion of the observed variance, except for *NPR1* (Table 4).

**Fig. 7.**
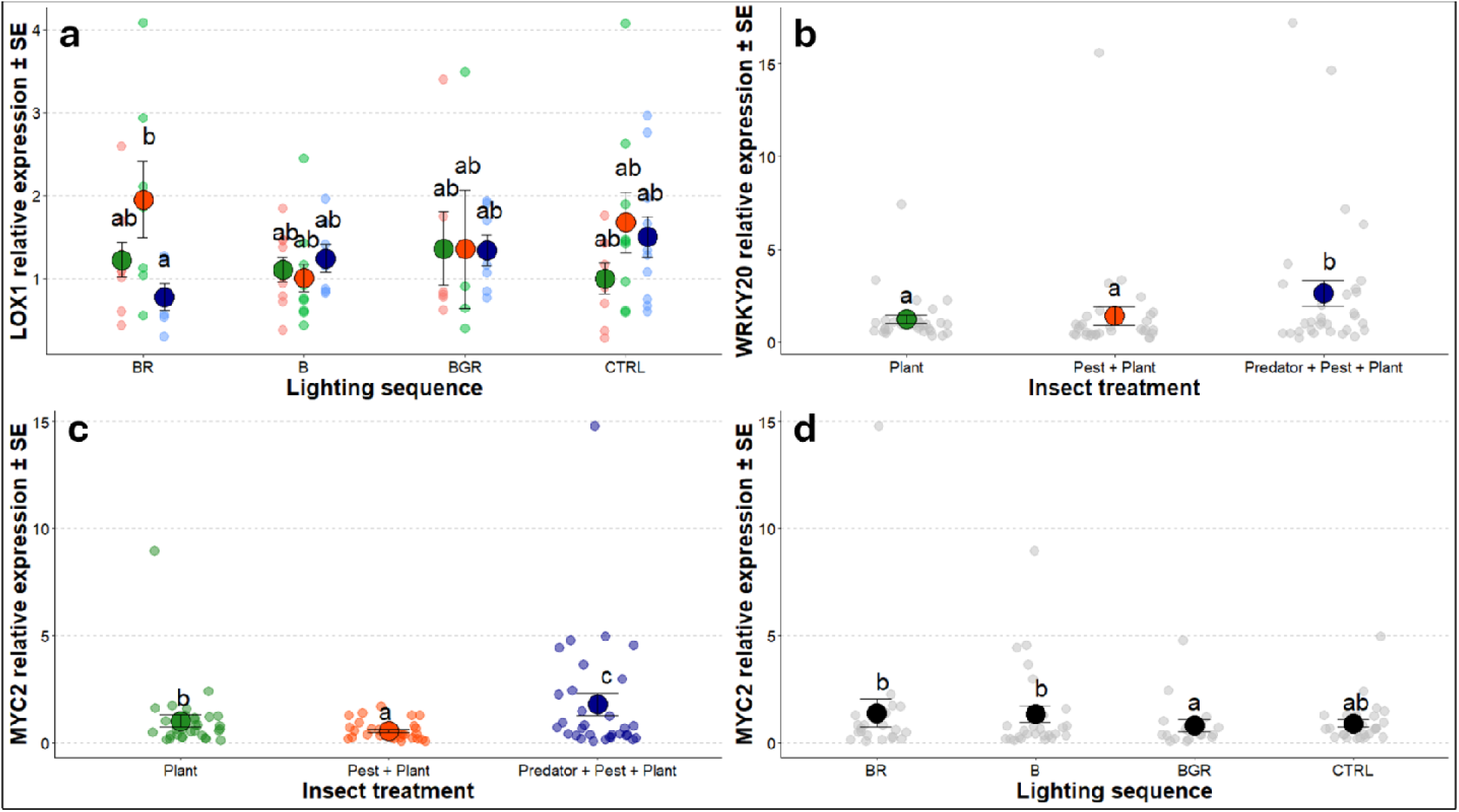
Jasmonic acid-related genes expression measured two weeks after exposing cucumber plants to different lighting spectra either in the absence of insects, with pests (*F. occidentalis*), or with pests and predators (*O. insidiosus*): a) *LOX1* expression (± SE) according to lighting sequence; b) *WRKY20* expression (± SE) according to insect treatment; c) *MYC2* expression (± SE) according to insect treatment; d) *MYC2* expression (± SE) according to lighting sequence. Pale points show raw data; darker points indicate model-predicted means. For each gene, expression is shown as fold change relative to the tested combination CTRL light × Plant-only

**Fig. 8.**
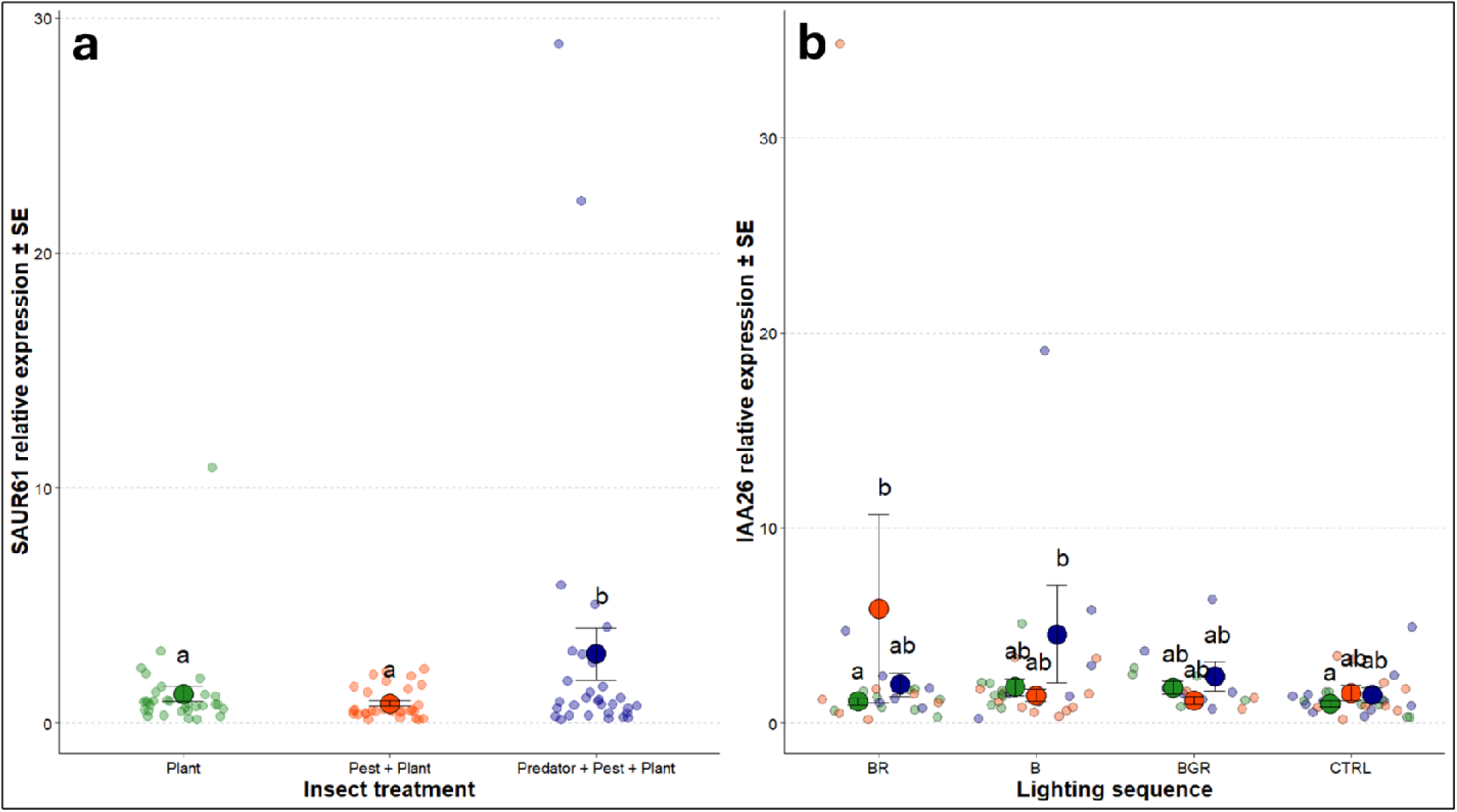
Auxin-related genes expression measured two weeks after exposing cucumber plants to different lighting spectra either in the absence of insects, with pests (*F. occidentalis*), or with pests and predators (*O. insidiosus*): a) *SAUR* expression (± SE) according to insect treatment; b) *IAA26* expression (± SE) according to lighting sequence. Pale points show raw data; darker points indicate model-predicted means. For each gene, expression is shown as fold change relative to the tested combination CTRL light × Plant-only

**Fig. 9.**
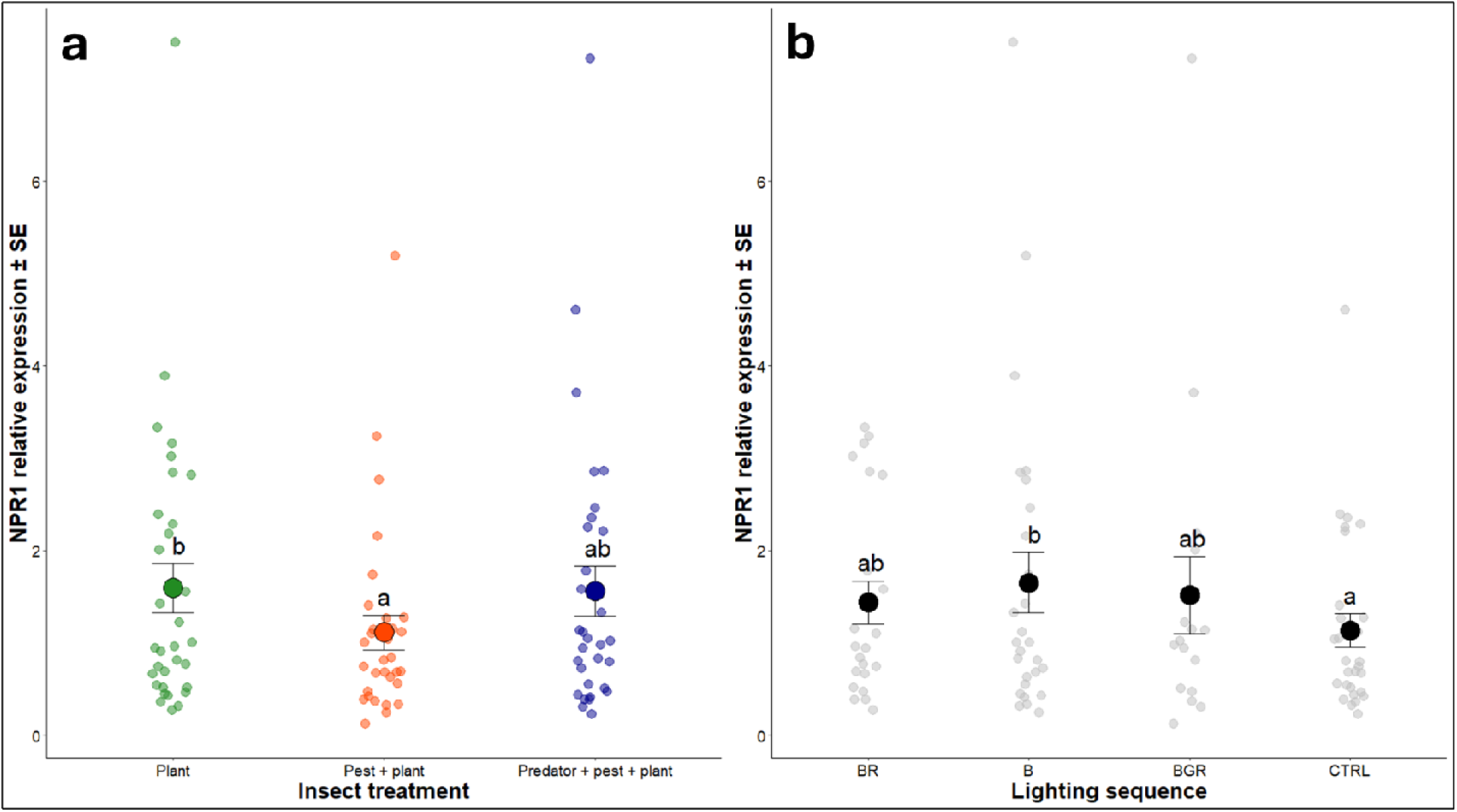
Salicilic acid-related gene expression measured two weeks after exposing cucumber plants to different lighting spectra either in the absence of insects, with pests (*F. occidentalis*), or with pests and predators (*O. insidiosus*): a) *NPR1* expression (± SE) according to insect treatment; b) *NPR1* expression (± SE) according to lighting sequence. Pale points show raw data; darker points indicate model-predicted means. For each gene, expression is shown as fold change relative to the tested combination CTRL light × Plant-only

**Table 4.**
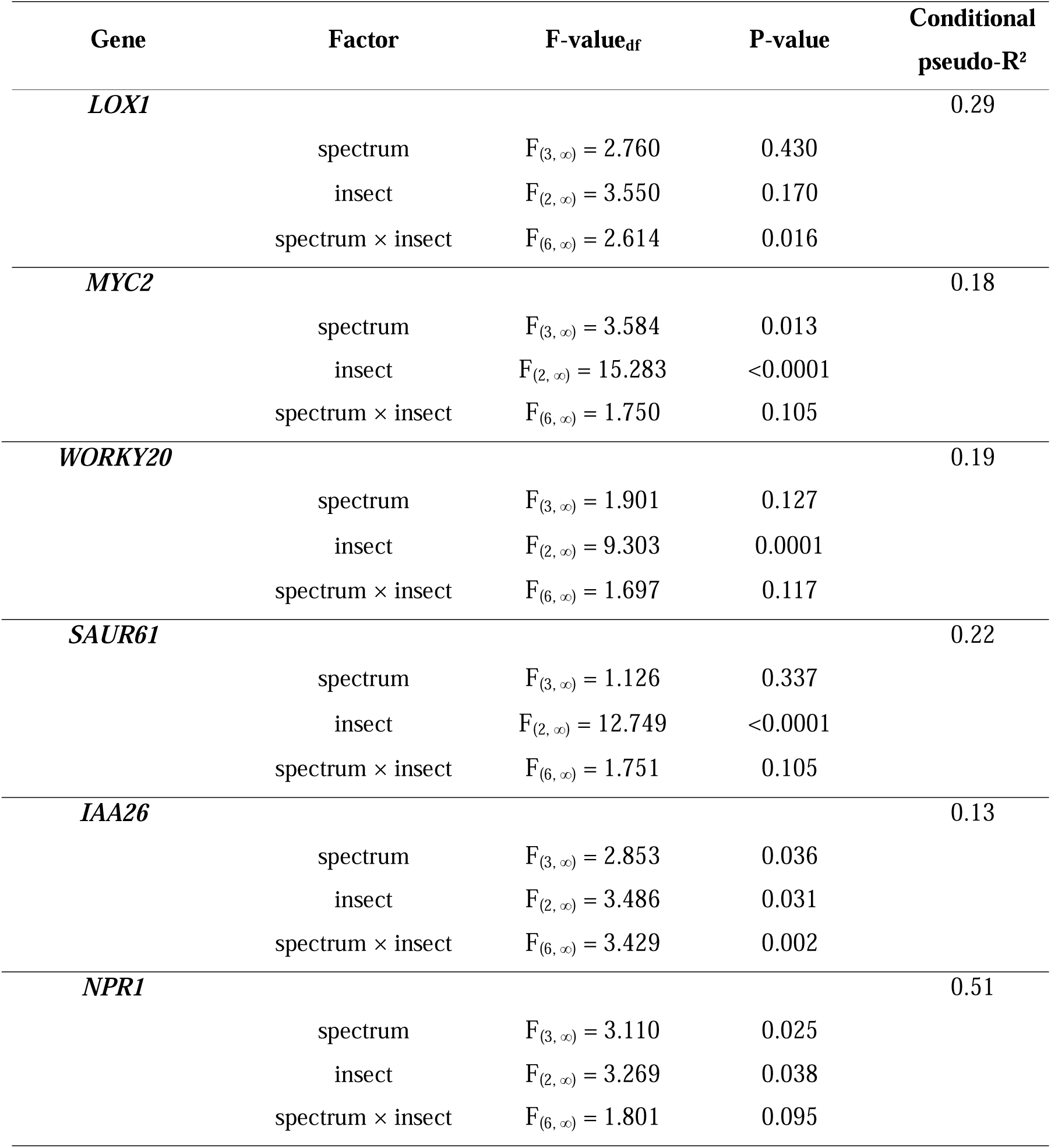
Statistical comparison of defense-related genes induction of plants exposed to different insect combinations and lighting sequences.

## Discussion

*Orius insidiosus* successfully established under all tested lighting conditions, consistently reducing thrips populations and limiting leaf damage, while the cucumber plants’ photosynthetic performance was not affected by the presence of pests or predators. Thrips densities in the pest-only treatment were similar across lighting conditions, indicating that light supplementation did not favor *F. occidentalis*. Notably, genes associated with the jasmonic acid (JA) pathway were unexpectedly upregulated in the presence of *O. insidiosus*, regardless of the light spectrum, suggesting that predator-related cues may influence plant defense signaling.

### Insect Densities and Silvering Damage

A consistent pattern of predator establishment was observed across all lighting sequences, with *O. insidiosus* populations tripling on average over the course of the experiment relative to the initial numbers released (25 ± 5; mean ± SE), although populations grew less rapidly under extended photoperiods with BGR spectra. Contrary to our initial prediction, predator abundance was not higher under blue light (Fig. 4A), and thrips suppression remained stable across all lighting treatments (Fig. 4B). Interestingly, although predator numbers were greater under the control condition (no photoperiod extension), their reduced abundance with extended photoperiod did not translate into higher pest densities (Fig. 4B) or increased silver damage (Fig. 4C), indicating that biological control efficacy remained unaffected by lighting regime. Although based on relatively short timeframes, these results may support the strategic use of specific light spectra for just a few days, rather than continuous exposure like we tested, to trigger key *O. insidiosus* behaviors, such as mating or prey searching, thereby promoting its establishment or predation (Canovas et al. 2025b). Using higher light intensities may also reveal contrasts between photoperiod-extension treatments for *O. insidiosus*- mediated pest control, as seen in less complex settings (Canovas et al. 2025a, b). Overall, predator presence—rather than differences in lighting—appeared to drive pest suppression (Fig. 4B), consistent with findings for other beneficials under LED lighting (Meijer et al. 2023; Athanasiadou et al. 2024; Fraser et al. 2024). These findings under controlled experimental conditions are also consistent with anecdotal observations from real production settings: *O. insidiosus* successfully established over winter in a commercial strawberry greenhouse under LED-supplemented natural light with BGR spectrum, achieving effective thrips control and reducing fruit damage compared to years without predator release (M.L. Canovas, pers. obs., Québec, January to May 2024). This growing body of evidence highlights the remarkable behavioral plasticity of biological control agents. However, longer-term studies—ideally spanning full cropping cycles—are still needed to assess whether varying artificial lighting in CEA can sustainably maintain *O. insidiosus* efficacy. Repeated exposures to specific spectra and recurrent predator introductions, as in commercial biocontrol programs, may also reveal cumulative plant protection benefits overlooked in shorter trials.

### Plant Morphological Traits and Light Assimilation

Plant morphological traits and light assimilation indices tended to be influenced by light treatments (Table 3) rather than the presence of pests and predators. As predicted, B and BR treatments led to the highest photosynthesis-related parameters (Fig. 6) and fresh biomass was significantly greater under B light, corresponding to the blue and red absorption peaks of chlorophyll a and b (Bantis et al. 2018). Dry biomass was also higher under LED supplementation including BGR wavelengths compared to no supplementation, as previously reported for pepper in a greenhouse context (Fraser et al., 2024). Photosynthetic performance, assessed via QY (Fv/Fm ratio), SPAD, and PI indices, was not affected by insect presence (Table 3), suggesting that silver damage caused by *F. occidentalis* did not impair photosynthetic capacity under the tested lighting conditions. Combined with similar pest densities observed across lighting sequences (Fig. 4B), this result indicates that the low-intensity light supplementation that we provided (∼30 µmol/m²/s) does not promote thrips proliferation, regardless of spectral composition. A comparable trend was reported for the aphid *Myzus persicae* (Sulzer) under ∼90 µmol/m²/s for various spectra (Fraser et al., 2024). These findings temper earlier concerns about the use of artificial lighting in commercial crop production, particularly the idea that extended photoperiod might enhance the activity or impact of diurnal pests by providing more nutritious hosts, leading to winter outbreaks (Vänninen et al. 2010). Yet, as lighting technologies evolve and production conditions diversify, it will still be important to assess the responses of key pest species to light regimes typical of greenhouses and indoor farming environments on a case-by-case basis.

### Plant defense and growth gene expression

Defense-related phytohormone induction varied across combined spectral and insect treatments. Unexpectedly, JA-pathway genes were mostly upregulated in predator treatments (Fig. 7), possibly reflecting a shift toward phytophagy by *O. insidiosus*— which is known to engage in some feeding on plants (Zeng and Cohen 2000) – as prey declined over time. The downstream signaling genes *MYC2* and *WRKY20* (Fig. 7B-C) showed stronger induction than the upstream biosynthetic gene *LOX1* (Fig. 7A) (Wasternack and Song 2017), except under BR light, where pest-only plants showed the most silvering damage (Supplementary material 2). Similar JA-associated gene expression have been reported in major crops after short-term exposure to *O. laevigatus* or *O. sauteri*, where predator feeding and oviposition reduced thrips performance and, in some cases, enhanced attraction of beneficials (De Puysseleyr et al. 2011; Bouagga et al. 2018; Zhu et al. 2024b). In another study on cucumber, *O. sauteri* pre-inoculation significantly reduced thrips reproduction (Di et al. 2022). Applying a similar pre-inoculation approach could clarify whether *O. insidiosus* triggers such effects, and whether light modulates this response, given its oviposition sensitivity to light (Canovas et al. 2025b). Transiently high insect densities in our predator treatment may also have boosted DAMP/HAMP signaling (Acevedo et al. 2015). However, hormone response amplitudes were slightly lower than those reported in previous *Orius* spp. studies, consistent with the moderate explanatory power of our models (Table 4) and were likely influenced by the timing of our sampling. Plant defense responses, including defense gene expression, fluctuate over time, allowing for fine-tuned adaptation to dynamic environmental cues. Unlike most studies measuring gene expression shortly after insect or light exposure (Abe et al. 2008; Mirzahosseini et al. 2020; Zhu et al. 2024b) we sampled two weeks later. This is the first report of JA activation in cucumber by *O. insidiosus* after prolonged exposure, independent of light conditions—a realistic scenario of plant–predator interactions under lighting regimes typical of CEA systems. Future studies using our lighting setup should investigate temporal dynamics to better capture defense induction peaks and enable direct comparisons with existing literature.

Auxin- and salicylic acid–related gene responses in our system were shaped by both insect presence and light treatments, although the responses were inconsistent, and the effect sizes were small. Contrary to our predictions, expression of the SA-associated gene *NPR1* was modestly upregulated under B light and in the absence of insects (Fig. 9), whereas BR lighting increased transcript levels of the AUX-related *IAA26* gene in BR-exposed, pest-only plants (Fig. 8B). Beyond the effects of sampling time mentioned earlier, our lighting treatments were milder than the extreme regimes often applied in plant physiology studies, which typicallyrely on high light intensities and narrowband artificial spectra to isolate hormonal processes (Vänninen et al. 2010; Pierik and Ballaré 2021; Lazzarin et al. 2021). More studies are needed to demonstrate how insect activity and lighting conditions interact under experimental setups that better reflect commercial CEA conditions.

### Summary and conclusions

Our results highlight the robustness of *O. insidiosus* as a biological control organism across different lighting conditions, over several weeks in a complex environment mimicking CEA systems. Consistent with previous studies under LED lighting, pest suppression was primarily driven by the presence of biological control organisms (top-down effects) rather than by lighting conditions. In our case, both predation and plant defense induction by predators may have contributed to reducing thrips populations and plant damage. Given the dual role of *Orius* spp. in exerting efficient direct predation and triggering plant defenses (Cox et al. 2006; Herrick et al. 2021; Bouagga et al. 2018; Zhu et al. 2024b), as well as their ability to reproduce and forage under various spectra (Wang et al. 2013; Labbé and McCreary 2020; Canovas et al. 2025a, b), these predators appear highly promising for use under artificial lighting in CEA. Further potential lies in integrating *Orius* spp. under intermittent or low-intensity LED lighting regimes, specifically designed to reduce energy consumption in CEA systems while enhancing the organoleptic quality of harvested crops (Gao et al. 2021; Boucher et al. 2023; Zou et al. 2025). As such, combining lighting strategies that jointly support plant physiology while maintaining beneficial insect performance could enhance the efficacy and resilience of pest management in CEA.

## Acknowledgments

The authors would like to acknowledge the partnership and financial support received from the Ministère de l’Agriculture, des Pêcheries et de l’Alimentation du Québec, MITACS, SOLLUM Technologies inc., ANATIS Bioprotection and the National Optics Institute (project n° IA120649 & IT17200-IT40569). The authors sincerely thank Virginie Simard (B.Sc., Biology) for her valuable technical assistance with the hormonal analyses.

## Authors’ contribution

MLC: Conceptualization; Funding acquisition; Project administration; Methodology; Investigation; Formal analysis; Visualization; Writing – original draft; Writing – review & editing. AB: Methodology; Investigation; Formal analysis; Writing – review & editing. PKA: Supervision; Writing – original draft; Writing – review & editing. JFC: Funding acquisition; Supervision; Writing – review & editing. TG: Funding acquisition; Supervision; Writing – review & editing. MD: Funding acquisition; Project administration; Supervision; Resources; Writing – review & editing.

---

**Supplementary information 1.**
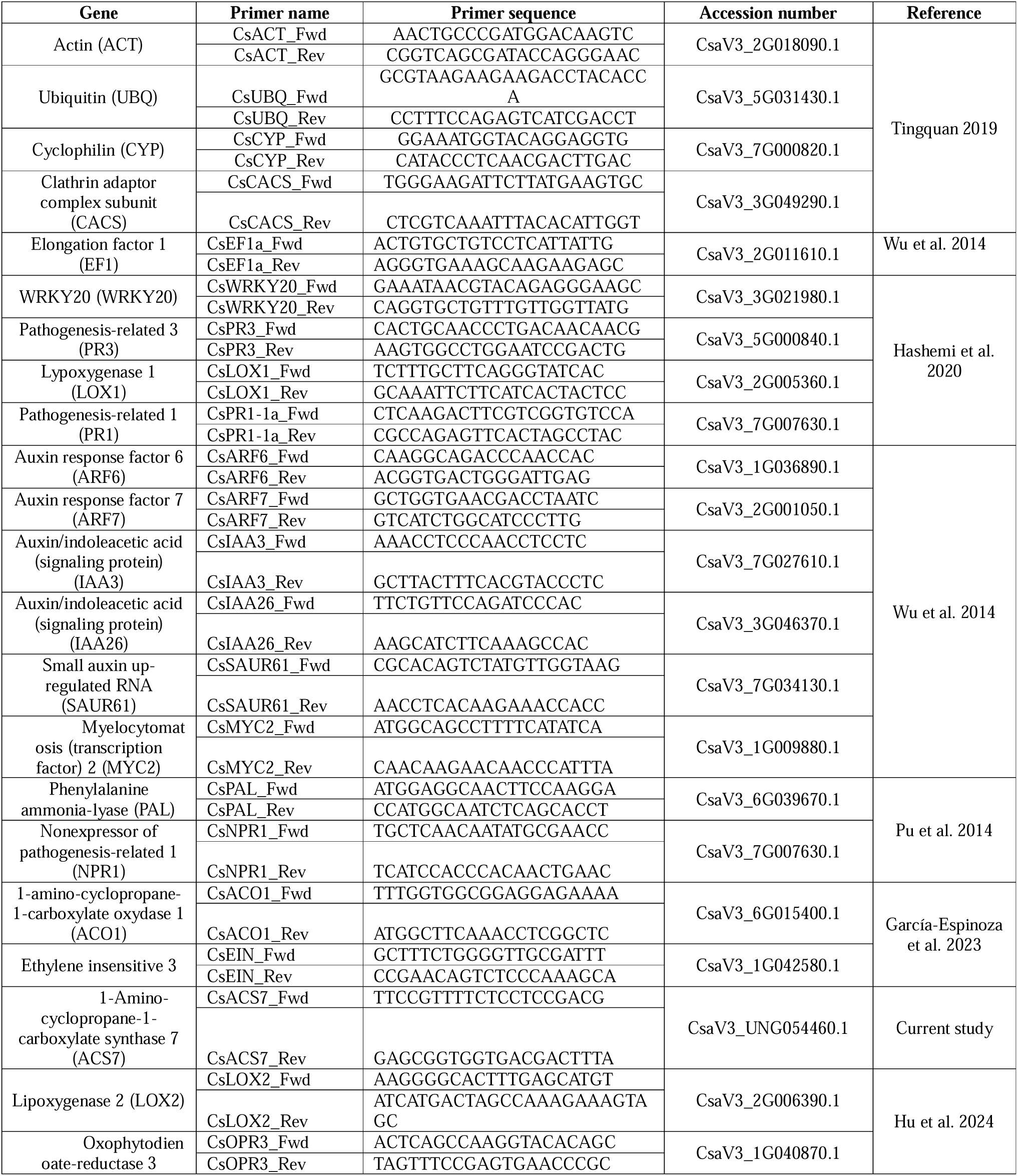
Primer sequences and associated references used for gene expression quantification by qPCR analysis.

**Supplementary information 2.**
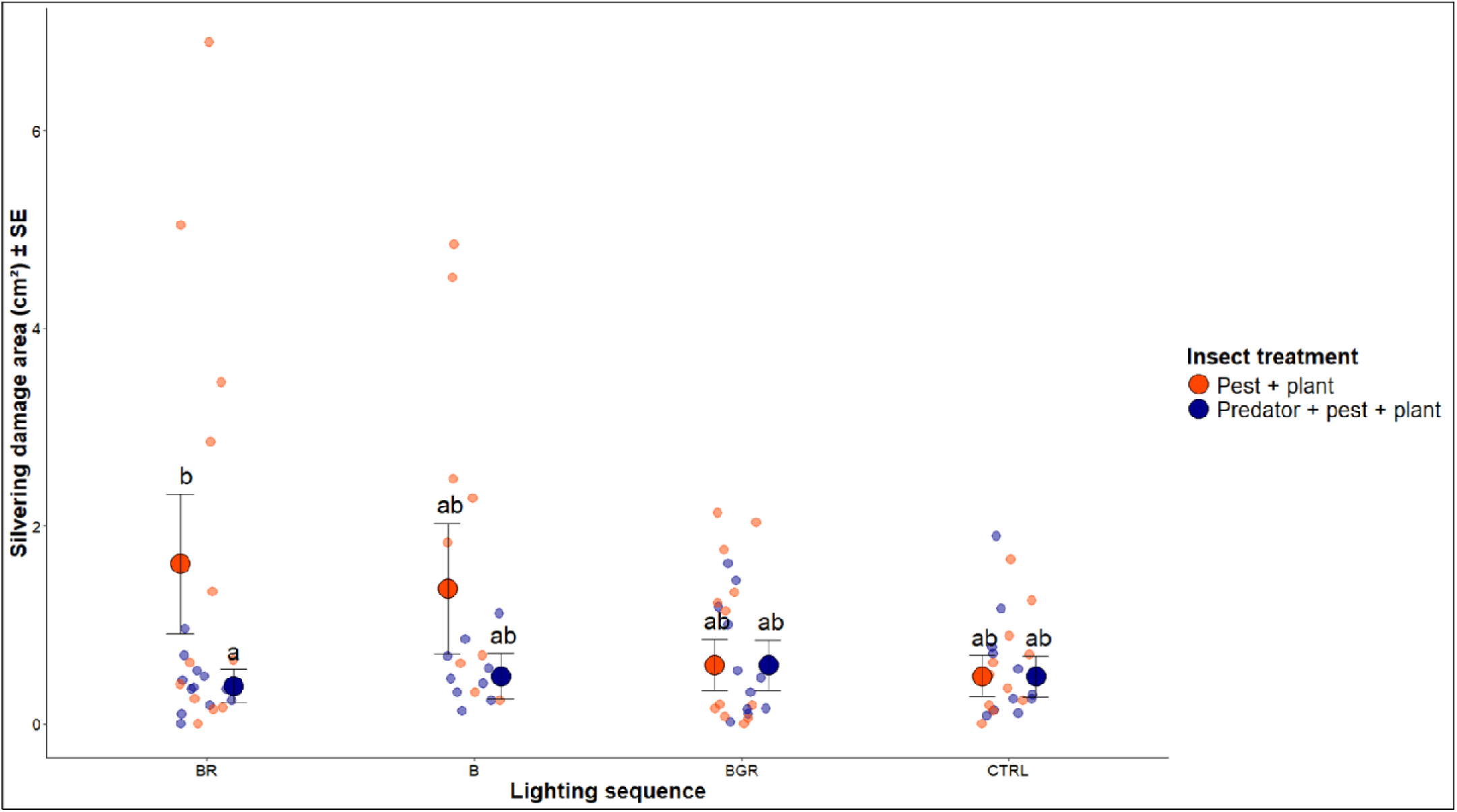
Silvering damage area in (cm²) (± SE) according to the interaction between lighting sequence and insect treatment. Pale points show raw data; darker points indicate model-predicted means.

**Supplementary information 3.**
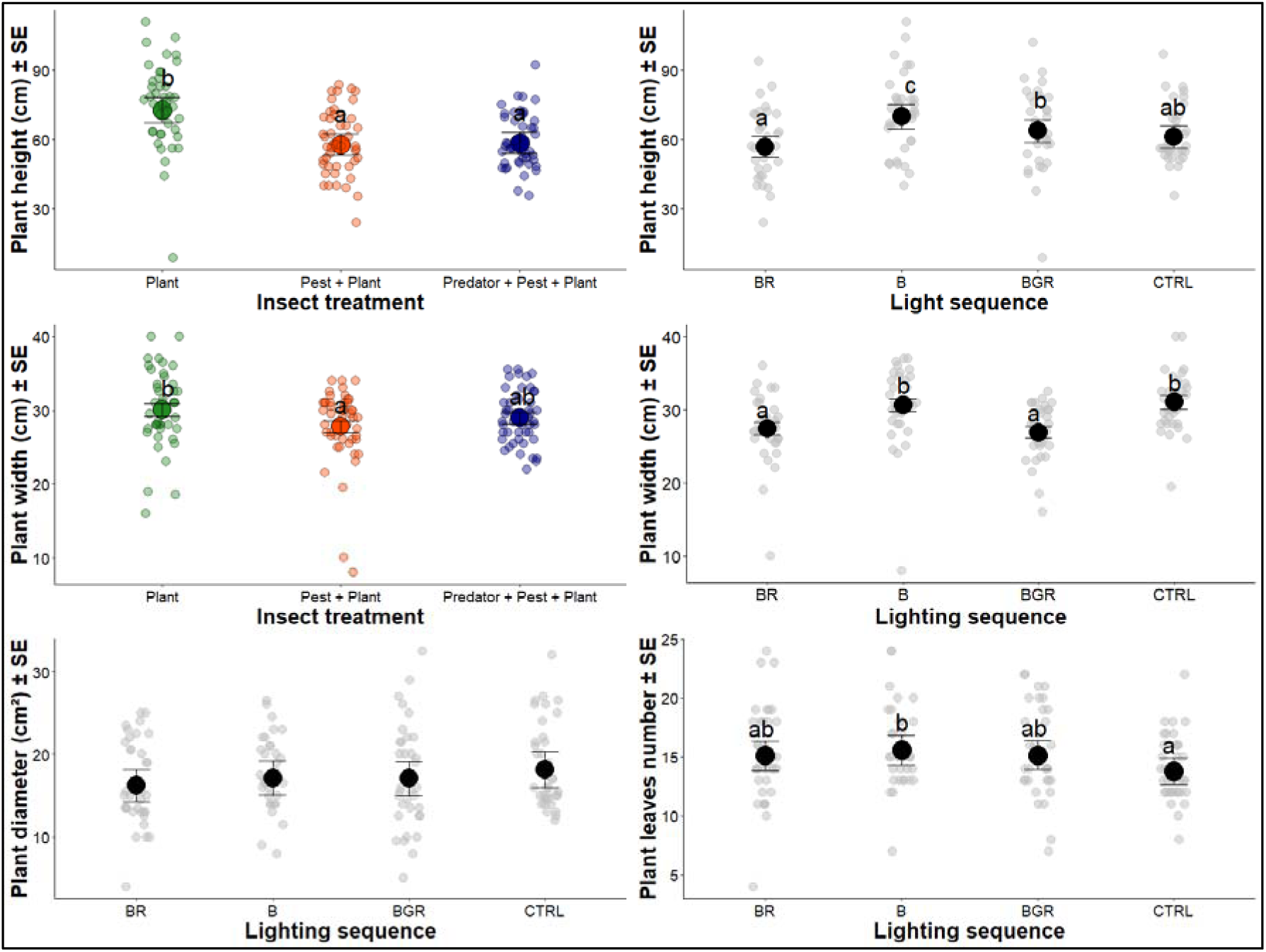
Additional plant morphological parameters measured during the cage experiment. Pale points show raw data; darker points indicate model-predicted means.

